# Biogeochemical and omic evidence for paradoxical methane production via multiple co-occurring mechanisms in aquatic ecosystems

**DOI:** 10.1101/2020.07.28.225276

**Authors:** Elisabet Perez-Coronel, J. Michael Beman

## Abstract

Aquatic ecosystems are globally significant sources of the greenhouse gas methane (CH_4_) to the atmosphere. However, CH_4_ is produced ‘paradoxically’ in oxygenated water via poorly understood mechanisms, fundamentally limiting our understanding of overall CH_4_ production. Here we resolve paradoxical CH_4_ production mechanisms through CH_4_ measurements, *δ*^13^CH_4_ analyses, 16S rRNA sequencing, and metagenomics/metatranscriptomics applied to freshwater incubation experiments with multiple time points and treatments (addition of a methanogenesis inhibitor, dark, high-light). We captured significant paradoxical CH_4_ production, as well as consistent metabolism of methylphosphonate by abundant bacteria—resembling observations from marine ecosystems. Metatranscriptomics and *δ*^13^CH_4_ analyses applied to experimental treatments identified an additional CH_4_ production mechanism associated with (bacterio)chlorophyll metabolism and photosynthesis by Cyanobacteria, and especially by Proteobacteria. Both mechanisms occured together within metagenome-assembled genomes, and appear widespread in freshwater. Our results indicate that multiple, co-occurring, and broadly-distributed bacterial groups and metabolic pathways produce CH_4_ in aquatic ecosystems.

## Introduction

Atmospheric concentrations of the potent greenhouse gas methane (CH_4_) have increased significantly due to anthropogenic activity, representing an important component of climate change (1). However, these increases are superimposed on substantial spatial and temporal variability in natural sources of CH_4_ to the atmosphere. Of all natural CH_4_ sources, freshwater lakes are particularly important but poorly understood, with their estimated contribution ranging from 6 to 16% of all natural CH_4_ emissions—despite accounting for only ~0.9% of the Earth’s surface area (2). CH_4_ emissions from lakes are conventionally viewed to be regulated by CH_4_ production (occurring predominantly in anoxic sediments) and subsequent CH_4_ oxidation in surface sediments and the water column (3). However, oversaturation of CH_4_ has been consistently observed in oxygenated waters of aquatic systems (4). This observation indicates that CH_4_ is produced under oxic conditions, and that the rate of CH_4_ production exceeds CH_4_ oxidation. Since archaeal methanogenesis is an obligate anaerobic process (5), oxic CH_4_ production is typically referred to as the “methane paradox,” and has been observed in oceans (6, 7), lakes (8, 9, 10), and aerobic wetland soils (11). Notably, paradoxical CH_4_ production occurs near the surface, and so any produced CH_4_ may readily flux to the atmosphere. Identifying which mechanisms produce CH_4_ in oxygenated waters is therefore essential for our understanding of CH_4_ fluxes and their contribution to climate change.

Although multiple mechanisms for paradoxical aerobic CH_4_ production have been proposed, the degree to which these are active in freshwater lakes remains unknown. Initial studies suggested that CH_4_ production under oxygenated conditions could be occurring in anoxic microsites in the water column—such as fecal pellets, detritus, and the gastrointestinal tracts of larger organisms such as fish or zooplankton (12, 13, 14). Several studies have also demonstrated a correlation between phytoplankton or primary production and CH_4_ production (8, 9, 10). However, the underlying reason(s) for this relationship is unknown. One possibility is that methanogens reside on the surface of phytoplankton cells and produce CH_4_ in presumably anoxic microsites (8). Bogard et al. (9) and Donis et al. (15) also hypothesized that several groups of methanogens have oxygen-tolerant or detoxifying pathways that could aid in CH_4_ production in the presence of oxygen. For example, Angle et al. (11) characterized a methanogen candidate that possesses the enzymes to detoxify oxygen and produce CH_4_ under aerobic conditions.

In contrast, the current prevailing view of marine ecosystems is that methylphosphonate (MPn) is the main precursor of CH_4_ production under oxic conditions—particularly in phosphorus (P)-stressed ecosystems such as the open ocean (6, 16). MPn is the simplest form of organic carbon (C)-P bonded compounds in aquatic ecosystems; microbial utilization of MPn, and the consequent breakdown of the C-P bond, releases CH_4_ as a by-product (6, 16, 17). A broad range of marine and freshwater bacteria have the genomic potential to metabolize MPn and produce CH_4_, based on the presence of the multi-gene C-P lyase pathway in their genomes. This includes multiple groups of Proteobacteria, Firmicutes, Bacteroidetes, Chloroflexi, and Cyanobacteria (17, 18, 19). While expression of this pathway is thought to be regulated by phosphate availability (17, 18, 19), the degree to which this occurs in freshwater ecosystems is not well known (20, 21, 22). Finally, recent work indicates that cultures of marine and freshwater Cyanobacteria can directly produce CH_4_ (23). However, outside of a single experiment (24), this has not been examined in aquatic ecosystems. More significantly, the exact mechanism by which this occurs remains unknown. Given the widespread distribution of cyanobacteria in the ocean and freshwater, identifying the potential mechanism(s) by which cyanobacteria produce methane—and whether this capability may be present in other photosynthetic organisms—is of broad relevance.

These proposed mechanisms for CH_4_ production—(1) methanogenesis aided by detoxifying genes or in anoxic microsites, (2) CH_4_ production by breakdown of methylated compounds, and (3) CH_4_ production by Cyanobacteria—point to multiple mechanisms by which CH_4_ can be produced under oxygenated conditions. Many of these are recently discovered and therefore poorly understood, and the degree to which they occur within different aquatic ecosystems is largely unknown. We developed an experimental approach to disentangle these mechanisms and determine which may produce CH_4_ in freshwater lakes. We conducted incubation experiments using surface waters from high-elevation lakes, in order to rule out physical transport and focus on potential biological mechanisms of oxic CH_4_ production. We investigated specific mechanisms using a combination of CH_4_ measurements over time, experimental treatments and inhibitors, and 16S rRNA gene and transcript sequencing, while also applying stable isotope analyses and metagenome and metatranscriptome sequencing to selected experiments. Paradoxical CH_4_ production was evident in multiple experiments and experimental treatments, and could be attributed to MPn breakdown via widely-distributed members of the *Comamonadaceae* family. However, experimental treatments, stable isotope *δ*^13^C signatures of CH_4_, and metatranscriptomic data also point to a new potential mechanism of aerobic CH_4_ production carried out by photosynthetic bacteria.

## Results and Discussion

Our experiments provide multiple lines of evidence for paradoxical CH_4_ production in freshwater lakes. Out of 19 total experiments conducted in five lakes in Yosemite National Park, 26% of experiments showed unequivocal, monotonic CH_4_ production in unamended controls; 16% showed net oxidation in controls; 21% exhibited significant nonlinear patterns; and at least one experimental treatment in 37% of experiments also showed significant production (see below; based on replicate bottles incubated for at least 24 hours; Fig. 1). We observed the highest CH_4_ production rates in Lukens Lake (e.g., L1 and L2 experiments), as well as consistent production in Lower Gaylor Lake (LG1, LG3, and LG4) and Upper Cathedral Lake (UC1, UC2, UC3, and UC4). Net CH_4_ production rates ranged from 0.98 to 22.8 nM/h, with the majority of values <4 nM/h. These rates are consistent with the limited experiments previously conducted in other freshwater lakes—e.g., 0.1-2.5 nM/h and 3.7 in Lake Stechlin, Germany (8 and 10 respectively)—and, on the higher end, our values are similar to the range reported by Bogard et al. (9) for experimental manipulations in Lac Cromwell (2.08-8.33 nM/h). CH_4_ turnover rates also ranged from 5 to 146 days (with in situ concentrations ranging from 309-2839 nM), consistent with turnover rates of ~18 days in Lake Stechlin (CH_4_ concentration ~430 nM; 8), ~2.2 days in Lac Cromwell (CH_4_ concentration ~200 nM; 9), and 67 days in Yellowstone Lake (CH_4_ concentration of 46.3 nM; 21). Given these consistencies with observed rates in other lakes, paradoxical CH_4_ production rates in Yosemite appear broadly relevant.

**Fig. 1.**
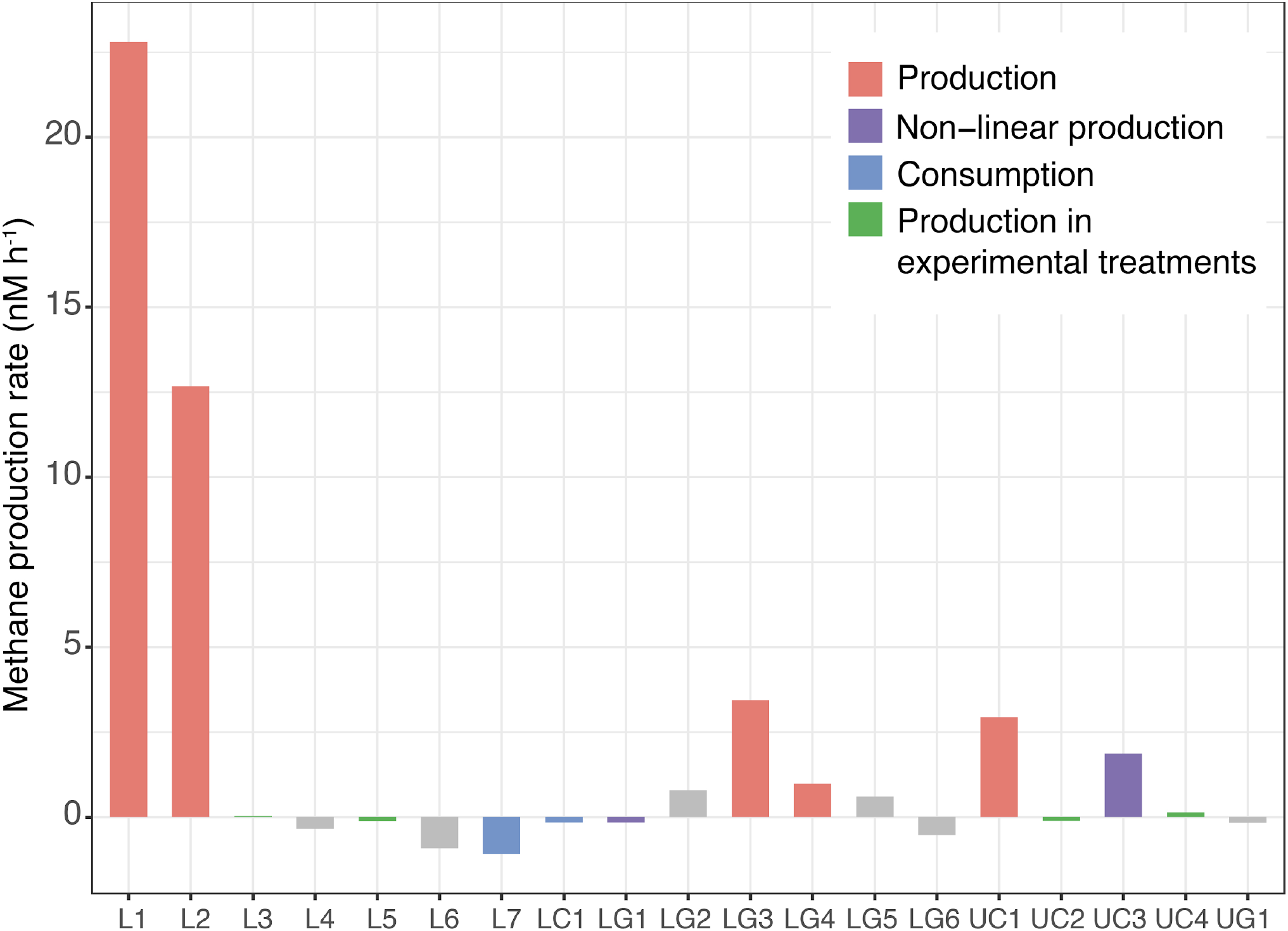
Net methane (CH_4_) production or consumption rates over 24 hours in unamended controls from each experiment. Different colors represent significant linear increases (production; red), linear decreases (consumption; blue), significant nonlinear patterns (purple), production in at least one treatment (green), or no significant change in CH_4_ concentrations (grey). Abbreviations denote lake and experiment number (e.g., UC4 is the fourth experiment in Upper Cathedral lake).

Eleven experiments did not show monotonic increases or decreases, which could reflect either (i) a complete absence of CH_4_ metabolism at the time of sampling, or, more likely, (ii) variations in the balance between co-occurring CH_4_ production and consumption over time. For example, initial CH_4_ production could be followed by oxidation once CH_4_ concentrations reach a certain threshold required for oxidation (8). Conversely, initial decreases due to oxidation may be followed by eventual production—for example through the development of P limitation that triggers CH_4_ production via MPn metabolism (6).We tested for nonlinear patterns using ANOVA, and found two cases of initial CH_4_ production followed by oxidation (LG1 and UC3; Figure S1 and SI Text). It is also possible that production and oxidation proceed at similar rates, leading to no net change in concentrations, even though CH_4_ is actively cycled. In this case, molecular data are useful for providing insight into the underlying dynamics. We consequently examined expression of the particulate methane monooxygenase gene *(pmoA)* in metatranscriptomes, and found that *pmoA* was expressed in the majority of the incubations surveyed (SI Text).

Observed decreases in CH_4_ concentrations, as well as omic data, are indicative of active CH_4_ oxidation that may obscure paradoxical production. Put another way, CH_4_ is clearly produced in incubations showing initial increases (UC3, LG1) or significant net increases (L1, L2, LG3, LG4 and UC1); however, it may also actively occur in experiments showing no significant change (or even consumption) over time, if CH_4_ oxidation rates are equal to or greater than production rates. We therefore evaluated potential paradoxical CH_4_ production mechanisms across experiments using multiple experimental treatments (BES, dark, high-light), and applied 16S rRNA sequencing, metagenomics, metatranscriptomics, and *δ*^13^C stable isotopic analyses to samples collected during these experiments.

### Methanogenesis

We used these multiple approaches to test for phytoplankton- or particle-based methanogenesis, as initial observations in freshwater lakes showed methanogens attached to phytoplankton (8), while other work has suggested CH_4_ production can occur in anoxic microsites on particles (12, 13, 14). In all experiments from 2017 and 2018, we included a treatment that consisted of the addition of the methanogenesis inhibitor BES (2-bromoethanesulphonate). While there are some caveats with its use (25), BES is widely used, including earlier tests of the CH_4_ paradox (8, 10). In our experiments, BES had an inhibitory effect on CH_4_ production in four experiments, but actually significantly increased CH_4_ production rates in four experiments (see below), and had no significant effect in the five remaining experiments (Fig. S2). In parallel, 16S rRNA sequencing of DNA from experiments L1-6 and LG1-4 recovered no methanogen 16S sequences (out of 5.4 million total sequences; Table 1). Sequencing of 16S rRNA transcripts in RNA samples from experiments L5-6, LG5-6, and UC3-4 recovered methanogen sequences in only two samples (dark treatments from L5 and LG5), and total numbers were only 0.014% and 0.008% of all 16S rRNA sequences (Table 1). Analysis of metatranscriptomes likewise showed that methyl coenzyme A (*mcrA*; responsible for methanogenesis) transcripts were absent in LG6 and UC4(tf incubations and present at low levels in LG5, L7 and UC4(t0) (Fig. 2a). *mcrA* genes were absent in all metagenomes, with the sole exception of the L6(tf) incubation (Fig. 2b). (Incubations were monitored to confirm that they were under oxic conditions at all times.) Collectively these data provide limited evidence—but not the complete absence of evidence—for water column-based methanogenesis as source of CH_4_, as BES reduced production in four experiments (Fig. S2), methanogens were almost entirely absent and inactive based on 16S data (Table 1), and *mcrA* transcripts and genes were rarely expressed and detected in metatranscriptomes and metagenomes (Fig. 2).

**Fig. 2.**
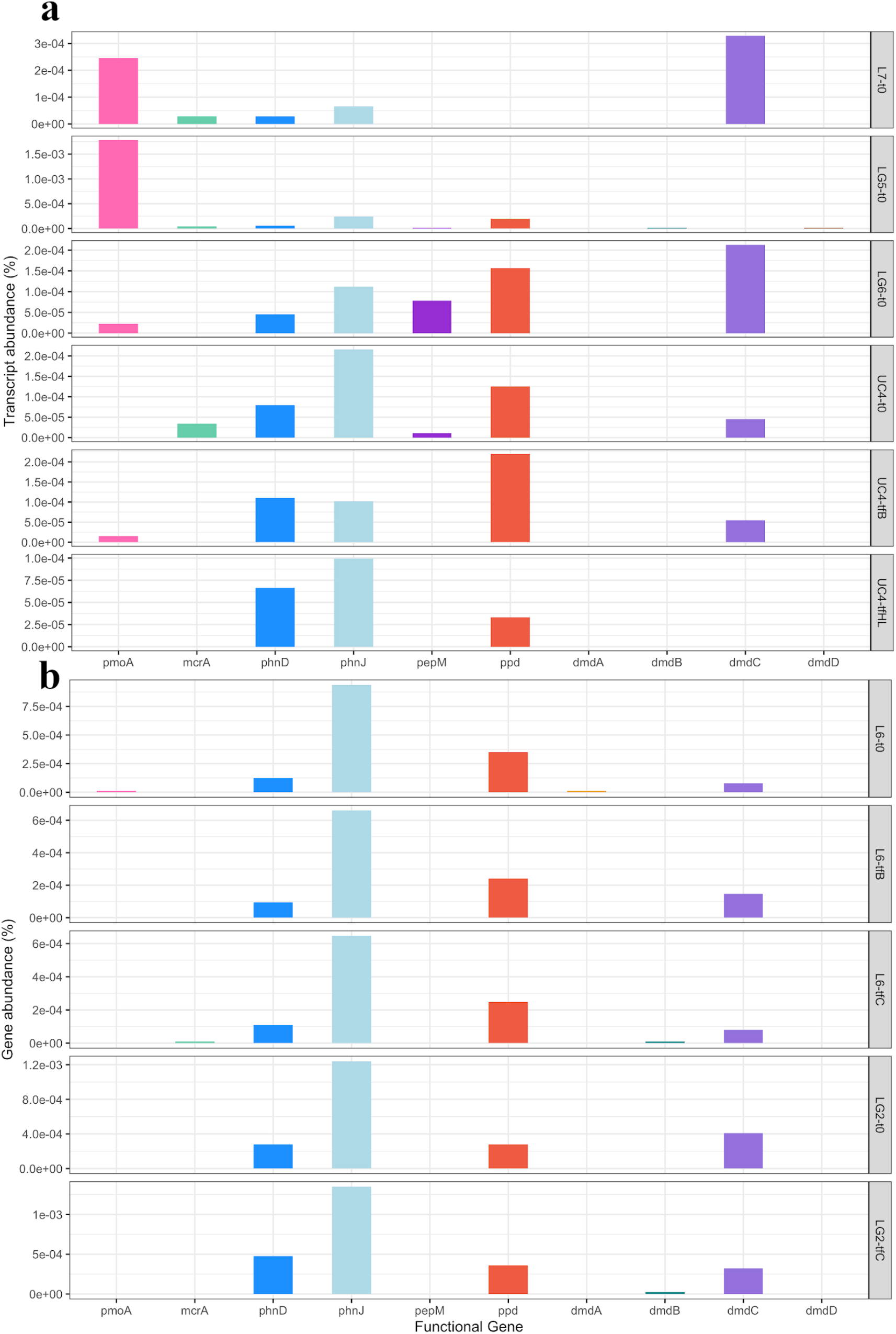
Variations across experiments and treatments in (a) transcript abundance (% of total reads of metatranscriptomes) and (b) gene abundance (% of total reads of metagenomes) of key functional genes. Functional genes quantified included those involved in: methane oxidation *(pmoA),* methanogenesis (*mcrA*), phosphonate assimilation *(phnD, phnJ)* and production *(ppd, pepM),* and DMSP metabolism *(dmdA, dmdB, dmdC, dmdD).*

**Table 1.** Relative abundance of methanogens, Cyanobacteria, and *Comamonadaceae* as a % of 16S rRNA genes and transcripts sequenced in different experiments and treatments

| Exp. | Treatment | Time (h) | Methanogens | Cyanobacteria | *Comamonadaceae* |
| --- | --- | --- | --- | --- | --- |
| % of 16S rRNA genes |  |  |  |  |  |
| L1 | Control | 0 | 0 | 0.902 | 9.17 |
| L2 | Control | 0 | 0 | 1.10 | 6.03 |
|  | Control | 24 | 0 | 1.65 | 5.20 |
| L3 | Control | 0 | 0 | 0.061 | 4.56 |
| L4 | Control | 0 | 0 | 2.38 | 7.97 |
| L6 | Control | 0 | 0 | 5.55 | 2.03 |
|  | BES | 45 | 0 | 12.6 | 0.338 |
| LG1 | Control | 0 | 0 | 0.646 | 5.32 |
|  | Control | 24 | 0 | 0.377 | 2.04 |
| LG2 | Control | 0 | 0 | 0.795 | 11.1 |
| LG3 | Control | 24 | 0 | 0.246 | 0.881 |
| LG4 | Control | 0 | 0 | 5.79 | 4.35 |
| UC1 | Control | 0 | 0 | 0.880 | 11.6 |
| % of 16S rRNA transcripts |  |  |  |  |  |
| L5 | Control | 0 | 0 | 46.0 | 6.43 |
|  | Control | 74 | 0 | 28.8 | 6.66 |
|  | Dark | 74 | 0.014 | 58.5 | 4.70 |
|  | BES | 74 | 0 | 24.9 | 8.65 |
| L6 | Control | 0 | 0 | 8.34 | 6.27 |
|  | Control | 55 | 0 | 0.800 | 5.42 |
|  | BES | 55 | 0 | 0 | 5.20 |
| L7 | Dark | 96 | 0 | 65.5 | 0.841 |
|  | BES | 96 | 0 | 46.4 | 1.95 |
|  | High light | 96 | 0 | 45.0 | 1.06 |
| LG5 | Control | 0 | 0 | 17.3 | 12.8 |
|  | Control | 86 | 0 | 2.00 | 12.0 |
|  | Dark | 86 | 0.008 | 19.2 | 10.2 |
|  | BES | 86 | 0 | 1.04 | 25.6 |
| LG6 | Control | 0 | 0 | 3.05 | 6.14 |
|  | Control | 96 | 0 | 2.45 | 8.69 |
|  | Dark | 96 | 0 | 2.65 | 2.53 |
|  | BES | 96 | 0 | 2.28 | 5.02 |
|  | High light | 96 | 0 | 7.18 | 2.64 |
| UC3 | Control | 0 | 0 | 0.227 | 7.98 |
|  | Control | 24 | 0 | 0.275 | 8.72 |
|  | Control | 48 | 0 | 0.221 | 15.2 |
|  | Control | 57 | 0 | 0 | 12.3 |
|  | Dark | 57 | 0 | 0 | 8.66 |
|  | BES | 57 | 0 | 0 | 10.9 |
| UC4 | Control | 96 | 0 | 3.148 | 14.0 |
|  | High light | 96 | 0 | 0.474 | 7.25 |

### Methylphosphonate metabolism as a source of methane

Instead, analysis of metatranscriptomes and metagenomes showed potential for CH_4_ production mechanisms other than conventional methanogenesis. In particular, we found evidence for microbial metabolism of MPn based on the universal expression of alpha-D-ribose 1-methylphosphonate 5-phosphate C-P lyase *(phnJ)* transcripts in metatranscriptomes (Fig. 2a). *phnJ* is responsible for the cleavage of the C-P bond that results in CH_4_ production from MPn; this mechanism is thought to be the dominant paradoxical CH_4_ production mechanism in the ocean (16), but has been documented in only a limited number of freshwater lakes (20, 21, 22). In addition to *phnJ,* phosphonate-binding periplasmic protein *(phnD)* genes (involved in the binding component of phosphonate uptake; 26) were also universally expressed (Fig. 2). *phnJ* and *phnD* genes were also recovered in all metagenomes (Fig. 2b).

The majority of *phnJ* and *phnD* genes were present in organisms of the same genera as the metatranscriptomes, and over 40 different bacterial genera expressed or possessed *phnJ* genes. This indicates that P assimilation from MPn is broadly distributed among surface water microbes in these lakes. However, *phnJ* transcripts and genes were predominantly found among organisms in the betaproteobacterial *Comamonadaceae* family: multiple *Comamonadaceae* genera accounted for 20.9% of *phnJ* transcripts in metatranscriptomes and 27.4% of *phnJ* genes in metagenomes (Table S1). *phnJ* transcripts and genes from *Sphingobacteriales* were also abundant in several samples (LG5 and LG6 metatranscriptomes, L6 metagenomes). *Comamonadaceae* transcripts and genes showed notably high identity to database sequences, with the majority of sequences showing >92% and up to 100% amino acid identity to *Acidovorax, Hydrogenophaga, Limnohabitans, Polaromonas, Rhodoferax,* and *Variovorax phnJ* sequences (Table S1).

16S data are consistent with this and demonstrate that *Comamonadaceae* were universally present and abundant across experiments. In 16S rRNA gene sequence libraries from experiments L1-6, LG1-4, and UC1, *Comamonadaceae* ranged from 2.04% (L6) to 11.6% (UC1) of all sequences (Table 1), and were abundant in several incubations with significant CH_4_ production (L1, L2, UC1). Sequencing of 16S rRNA transcripts in RNA samples from experiments L5-6, LG5-6, and UC3-4 showed similar values (Table 1), and *Comamonadaceae* reached 15% of all 16S rRNA transcripts in the UC3 and UC4 experiments (where CH_4_ was significantly produced in some treatments). Similar to *phnJ* results, 16S sequences from the dominant *Comamonadaceae* ASVs showed high (98.8-100%) nucleotide sequence identity to 16S rRNA genes in sequenced genomes (Table S2).

Finally, we assembled and annotated metatranscriptomes and metagenomes to examine whether particular groups possess and express multiple *phn* and related genes in parallel, as the genes involved in phosphonate acquisition and utilization often cluster in genomes (18). We found that *phnJ* genes commonly clustered with other *phn* genes on the same contigs, particularly those from the *Comamonadaceae* (Table S3). For example, contigs >50 kb containing *phnC, phnD, phnE, phnI, phnJ, phnK, phnL, phnM* were present in metagenomes from the LG2 and L6 incubations, and were affiliated with the abundant freshwater bacterial group *Limnohabitans* (31, 32, 33), as well as other members of the *Comamonadaceae,* such as *Hydrogenophaga* and *Acidovorax.* Several longer contigs also contained *pst* or *pho* genes involved in P uptake and metabolism (Table S3). P limitation is common in freshwater lakes, as it is in the open ocean (34, 35, 36). High-elevation lakes in the Sierra Nevada are typically oligotrophic (35, 37), and inorganic phosphate concentrations in the lakes sampled here are consistently near the limits of detection (100 nM) by colorimetric techniques, with the vast majority of measurements <200 nM (38, 39). Under these conditions, organic P compounds— including phosphonates—could serve as important sources of P. Our findings are consistent with this idea, and the presence and expression of *phnJ* genes indicates that several microbial groups are capable of releasing CH_4_ through the cleavage of the C-P bond in MPn (Fig. 2). Our data in particular implicate multiple genera within the *Comamonadaceae* in MPn metabolism, given high*phnJ* and 16S identities (Tables S1 and S2), recovery of contigs containing *phn* and other genes involved in P acquisition and metabolism (Table S3), and their widespread presence and abundance in all experiments (Table 1).

### Evidence for methane production by photosynthetic bacteria

In addition to microbial use of MPn, Cyanobacterial photosynthesis was recently identified as a possible source of paradoxical CH_4_ production—yet the pathway by which Cyanobacteria ultimately produce CH_4_ remains unknown (23). Based on 16S rRNA genes and transcripts, and metagenomes and metatranscriptomes, Cyanobacteria were common in our samples (Table 1). We used two additional treatments to partition the relative effects of CH_4_ production vs. oxidation and evaluate Cyanobacteria as a potential source of CH_4_. First, we conducted light versus dark treatments in some experiments, as (i) light has been shown to inhibit CH_4_ oxidation (40, 41, 42), and (ii) CH_4_ production in cyanobacterial cultures was positively correlated with light (23). As a result, dark treatments would be expected to have higher CH_4_ oxidation rates, lower cyanobacterial CH_4_ production, and therefore lower CH_4_ concentrations compared with controls. Additionally, we increased light intensity in four experiments, which may have a twofold effect: (i) greater inhibition of CH_4_ oxidation, and (ii) increased rates of CH_4_ production from Cyanobacteria or other phytoplankton (8, 9, 23). For both of these reasons, we expected to observe higher CH_4_ production rates under higher light levels.

Dark bottles showed mixed results, with four cases of CH_4_ consumption (L4, LC1, and LG6 at 24 hours and LG5 at 86 hours; all p<0.05), production in two cases (UC2 at 24 hours and L5 at 74 hours; both p<0.05), and no significant difference in four experiments (L3, L7, LG4 and UC3). In two of four experiments with high light treatments, we observed increased CH_4_ production (L2 and UC4; both p<0.05). The UC4 experiment was particularly illuminating, as higher light intensity significantly boosted CH_4_ production rates by 62-fold compared with the control (up to 3.2 nM/h; p<0.0005). This was among the strongest treatment effects observed across all experiments (Fig. S2). BES also increased CH_4_ production rate by 49-fold compared with control in the UC4 experiment (up to 2.5 nM/h; p<0.0005, Fig. S2). This counterintuitive effect suggests another source of CH_4_.

To examine these responses in greater detail, we analyzed both the δ^13^C stable isotope composition of CH_4_ produced during the UC4 experiment, as well as changes in gene expression in response to experimental treatments in metatranscriptomes. We also directly compared these results with the LG6 and L7 experiments, which did not show CH_4_ production in BES or highlight treatments. Consistent with a lack of significant CH_4_ production in these experiments, δ^13^CH_4_ from multiple sampling time points showed no significant variation between treatments in the LG6 and L7 experiments (Fig. 3a).

**Fig. 3.**
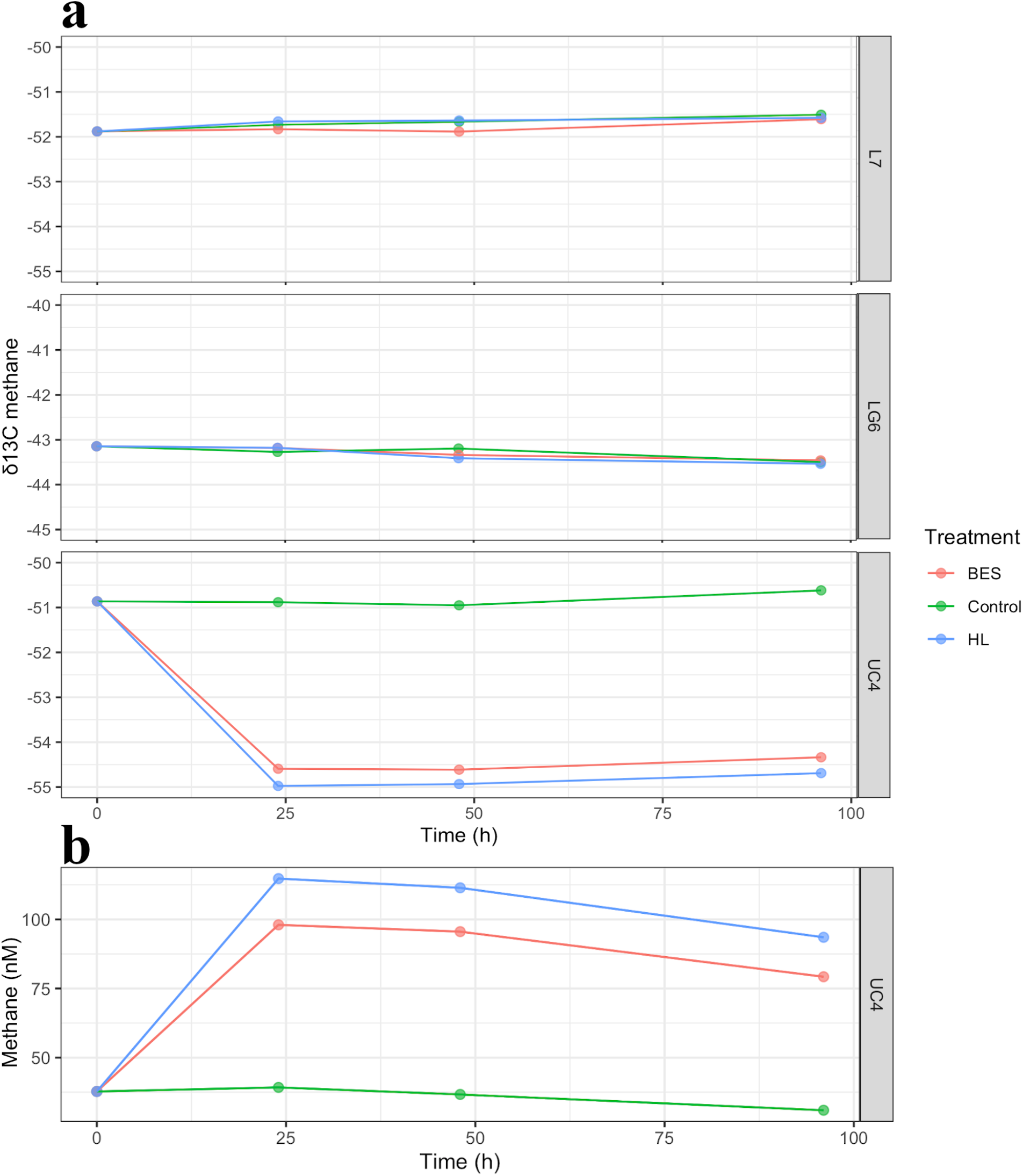
CH_4_ (a) δ^13^C values over time in the L7, LG6, and UC4 experiments and (b) concentrations over time in UC4 experiment. Different colors denote different experimental treatments. Scales of the vertical axes differ between experiments in panel a.

UC4 experimental data presented a clear contrast, as δ^13^CH_4_ values were significantly lower in the BES and high light treatments compared with controls (Fig. 3a). Along with increased CH_4_ concentrations (Fig. 3b), these data are indicative of an isotopically-depleted CH_4_ source in the BES and high light treatments, which we calculate to be −56.9‰ to −57.0‰ based on isotopic mass balance. These depleted values are substantially lower than atmospheric CH_4_ samples collected at the same time (−45.2 to −47.4‰), and reflect isotopic fractionation by the process producing CH_4_ (43). Methanogenesis does lead to strong fractionation, and our measured values fall at the upper end of typical δ^13^CH_4_ values for methanogenesis (−110 to – 50‰; 43). However, *mcrA* expression declined to undetectable levels (Fig. 2) in these treatments—which include BES, a methanogenesis inhibitor—suggesting another source. Strong isotopic fractionation from MPn is unlikely, as the mean isotopic fractionation for CH_4_ derived from MPn is only 1.3‰ (44). MPn could be allochthonous (terrestrial) or autochthonous (based on *pepM* results; Fig. 2), but in either case, is unlikely to be sufficiently ^13^C-depleted to produce the measured values. For example, the δ^13^C value of organic matter in surface waters of nearby Swamp Lake reaches a minimum of −37‰ (45). *phnJ* was also downregulated in both the BES and high-light treatments (Fig. 4), further suggesting that MPn breakdown is not a major source of CH_4_ in these treatments. However, our isotopic data and metatranscriptomic data are consistent with limited information available regarding photosynthetic CH_4_ production, which indicates that this is an isotopically light source (−55 to −45‰; ref. 23, 46).

**Fig. 4.**
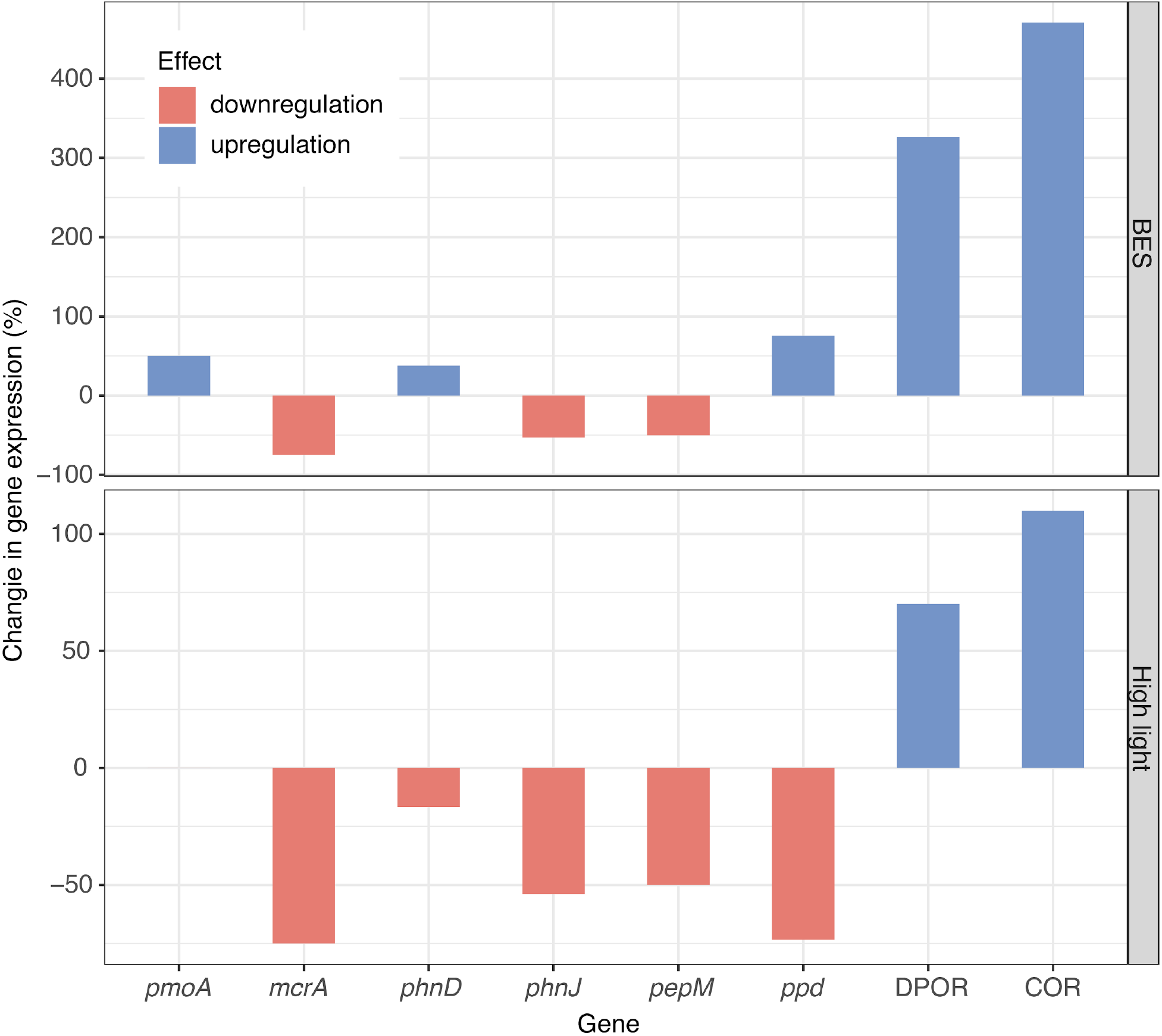
Upregulation and downregulation of genes involved in methane oxidation *(pmoA),* methanogenesis *(mcrA),* phosphonate assimilation *(phnD, phnJ)* and synthesis *(pepM, ppd),* and porphyrin and chlorophyll metabolism (DPOR and COR) in the UC4 experiment. BES treatment is shown in the top panel and the high-light treatment in the bottom panel. Colors denote the net effect of the treatment on the different genes in metatranscriptomes.

Although the pathway by which Cyanobacteria produce CH_4_ remains unidentified, Bižić et al. (23) hypothesized that CH_4_ is produced during photosynthesis owing to positive correlation between light and CH_4_. We used metatranscriptomic data to identify specific functions related to photosynthesis that were significantly different in the treatments vs. the control, and that may explain differences in CH_4_ concentrations and isotopic composition. This is an advantage of ‘omic data, as it is possible to examine previously unidentified potential mechanisms for CH_4_ production. Through comparison of all functions across metatranscriptomes from the UC4 experiment, we identified two functions related to chlorophyll biosynthesis that were upregulated and could potentially explain the increases in CH_4_: ferredoxin:protochlorophyllide reductase (DPOR) and chlorophyllide a reductase (COR) are involved in porphyrin and chlorophyll metabolism, and both were significantly different between the control and the BES (p<0.0005) and HL (p<0.005) treatments (Fig. 4). While not directly involved in photosynthesis, DPOR is found in photosynthetic bacteria, Cyanobacteria, and green algae, and is involved in the light-independent reduction of protochlorophyllide (47, 48). COR catalyzes the first step in the conversion of chlorin to a bacteriochlorin ring during bacteriochlorophyll biosynthesis (49, 50, 51). Notably, both DPOR and COR are nitrogenase-like enzymes, and both were abundant in all metatranscriptomes and metagenomes (Fig. 5). DPOR was found in several cyanobacterial groups—such as *Pseudanabaena, Dolichospermum, Snowella* and *Oscillatoria*—in the metatranscriptome data (Table S1). However, both DPOR and COR were particularly common within the *Limnohabitans* and *Polynucleobacter* genera (36.3-54.5% of DPOR and COR transcripts; Table S1), as some strains of these abundant freshwater bacteria are capable of aerobic anoxygenic photosynthesis (33, 52).

**Fig. 5.**
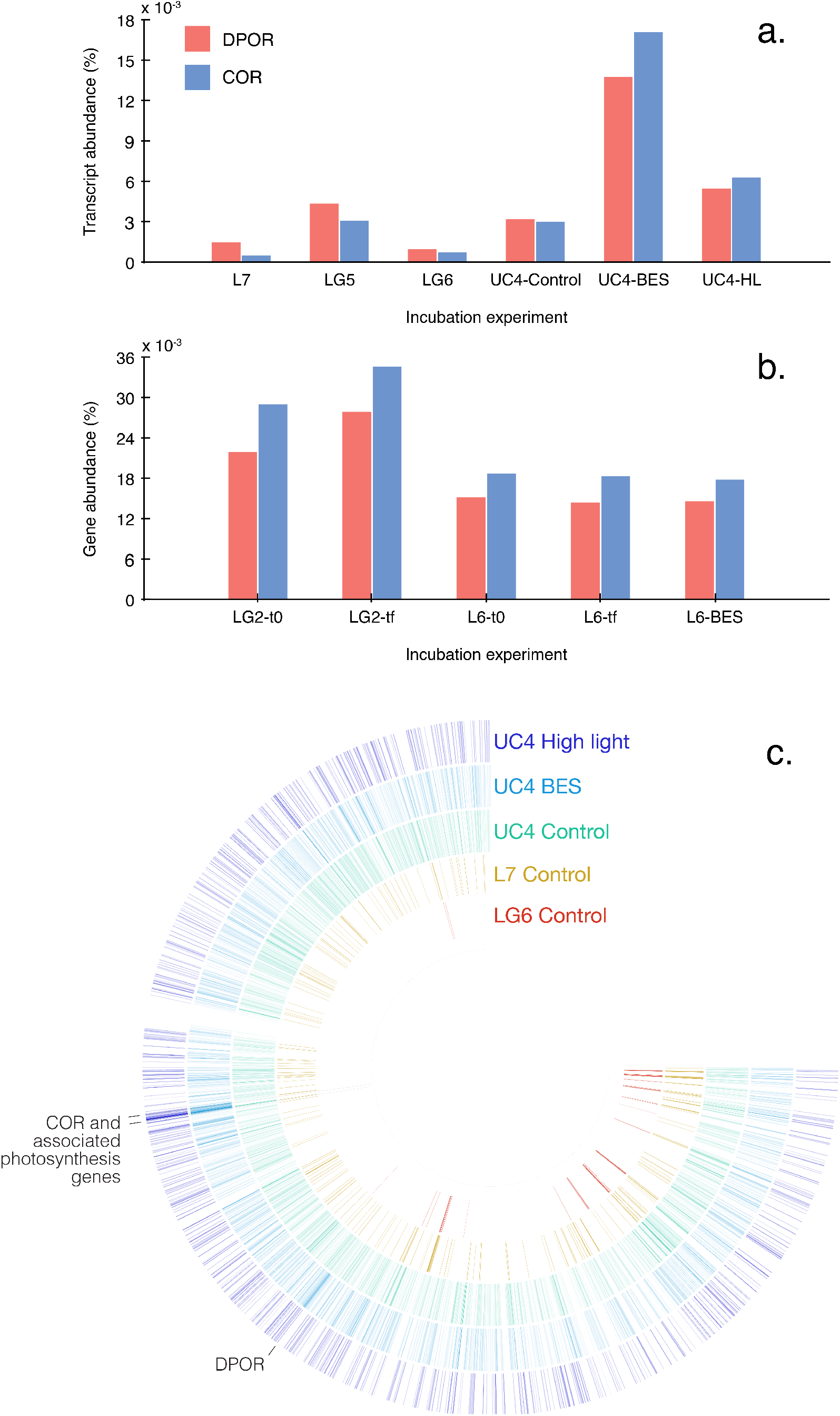
DPOR and COR reads in (a) metatranscriptomes, (b) metagenomes, and (c) mapped to the *Limnohabitans* metagenome-assembled genome (MAG) 5. In panels a and b, DPOR (red) and COR (blue) transcripts (a) and genes (b) are shown within different experiments and experimental treatments. In panel c, the *Limnohabitans* MAG 5 is shown in circular form, and the number of reads mapping to genes within the MAG are indicated by color intensity. The LG6 experiment is shown in red, L7 in orange, UC4 control in aqua, UC4 BES in light blue, and UC4 high-light in dark blue. Locations of DPOR and COR (and associated photosynthesis genes) within the MAG are indicated.

To further verify these findings, we mapped reads from the UC4 experiment, as well the LG6 and L7 experiments (which did not show CH_4_ production), to metagenome assembled genomes (MAGs). MAGs were assembled as part of the Genomes from Earth’s Microbiomes (GEM) catalog (53), with 27 MAGs recovered from two Yosemite metagenomes. Five of these MAGs were betaproteobacterial, including one *Rubrivivax* MAG, one *Polynucleobacter* MAG, and three *Limnohabitans* MAGs. All five of these betaproteobacterial MAGs contained genes for photosynthesis, including COR (Table S4). Three of these MAGs contained both COR and DPOR, as well as multiple genes involved in phosphonate metabolism. Of the remaining two MAGs, one lacked DPOR while the other lacked *phn* genes (Table S4), but these MAGs were estimated to be 55-58% complete.

Based on read mapping to these MAGs, we examined patterns in transcripts that showed differences in the BES and high light treatments of the UC4 experiment compared with the control. We also examined differences between the UC4 experiment versus the LG6 and L7 experiments. We found that multiple photosynthesis genes from *Limnohabitans* MAG 5 were highly expressed in UC4, while not detected at all in the LG6 and L7 experiments (Fig. 5c). Furthermore, of >12000 genes present in the five MAGs, several of these genes showed the highest overall number of mapped reads in the UC4 BES and high light treatments. Principal among these were two subunits of COR, as well as associated photosynthetic reaction center genes located on the same MAG scaffold (Fig. 5c). The *Limnohabitans* MAG 5 DPOR gene was also highly expressed. In contrast, phosphonate metabolism genes were expressed at lower levels and in similar levels in all metatranscriptomes.

DPOR and COR were expressed and present in all metatranscriptomes and metagenomes, yet the overall expression of DPOR and COR was an order of magnitude higher in the UC4 experiment compared with the LG6 and L7 experiments (Fig. 5a). Differences in the expression of DPOR and COR from *Limnohabitans* MAG 5 were even more pronounced. These patterns are highly consistent with the lack of a δ^13^CH_4_ isotopic signal in the LG6 and L7 experiments compared with the UC4 experiment (Fig. 3). Based on the positive effects of BES and of highlight on CH_4_ production observed in the UC4 experiments, this mechanism may also be active in the L2, L3, LG4, and UC3 experiments, where BES and high light also significantly increased CH_4_ production (Fig. S2). Decreases in CH_4_ in dark treatments in several experiments may also reflect decreased photosynthesis (alongside increased CH_4_ oxidation; Fig. S2). Experimental data also indicate that this mechanism is environmentally variable, given clear differences between experiments (Figs. 3 and 5). However, these variations are consistent with the fact that photosynthesis by *Limnohabitans* is variable, as photosynthesis is used to supplement heterotrophy (52).

Based on these findings, we propose two potential mechanisms that could be involved in paradoxical CH_4_ production in oxic surface waters of freshwater lakes: (1) CH_4_ may be produced from methoxyl groups present in (bacterio)chlorophyll precursors, or (2) CH_4_ production may be catalyzed by DPOR and COR enzymes. In terrestrial plants, CH_4_ is thought to be produced aerobically from structural components—such as pectin, lignin and cellulose (54, 55, 56)— mainly in plants under stress (such as increased temperature, UV radiation or physical damage; 57). Protochlorophyllide, chlorophyllide, and other bacteriochlorophyll precursors contain methoxyl groups that have been shown to serve as precursors of CH_4_ in plants (58), and a similar mechanism may be active in Cyanobacteria and/or Proteobacteria under stress (presumably due to BES additions and high light levels in our experiments).

Alternatively, CH_4_ production may be catalyzed by DPOR and COR enzymes. Nitrogenases reduce a range of multi-bond compounds (59, 60, 61, 62, 63), and this quality is shared across different nitrogenases (64). We propose that, similar to the findings of Zheng et al. (62), the nitrogenase-like enzymes DPOR and COR may be able to catalyze the reduction of CO2 into CH_4_ when a suitable source of electrons is present. This may explain why BES and high light increased CH_4_ production in some experiments, as BES could act as a source of electrons similar to thiosulfate in Zheng et al.’s experiments, while high light levels may disrupt photosynthesis and drive the buildup of electrons and reducing equivalents. Importantly, DPOR and COR are central in photosynthesis, and so are present in Cyanobacteria, as well as within the abundant and ubiquitous freshwater bacterial groups *Limnohabitans* and *Polynucleobacter* (31, 33, 52).

### Multiple co-occurring mechanisms produce methane in aquatic ecosystems

Our combined results indicate that multiple paradoxical CH_4_ production mechanisms are active in freshwater ecosystems, and have several implications for our understanding of aquatic CH_4_ production. First, confirmation of the CH_4_ paradox in freshwater is still limited in scope and relatively recent (8, 9, 23), yet freshwater lakes are potentially large sources of CH_4_ to the atmosphere (2). We conducted multiple experiments in multiple lakes, and measured significant CH_4_ production in controls and/or treatments in 68% of experiments (Fig. 1). In many of the remaining experiments, variations in CH_4_ concentrations over time, across experimental treatments, and in *pmoA* gene expression patterns suggest that CH_4_ production occurs in parallel with coupled CH_4_ oxidation (Fig. 2, SI Text). Because CH_4_ oxidation may obscure CH_4_ production, use of different experimental treatments combined with ‘omic data is required to identify paradoxical CH_4_ production. Our findings therefore expand on limited data and demonstrate that paradoxical CH_4_ production is common, but variable, in freshwater.

Given the ubiquity of *phnJ* genes in our data (Fig. 2), their presence on contigs from abundant organisms (Tables S1-S3), and the potential for P limitation, microbial metabolism of MPn may represent an important baseline CH_4_ production mechanism in freshwater lakes. This fits with current understanding of marine ecosystems (6, 16). Metatranscriptomic and metagenomic data also indicate that, like the ocean, DMSP transformations may be active in freshwater—with potential implications for both DMS and CH_4_ production. Our data further demonstrate that MPn may itself be produced within lakes based on *ppd* and *pepM* expression (Fig. 2). Quantifying the relative contributions of allochthonous vs. autochthonous MPn production—as well as broader P acquisition dynamics—is therefore essential to understanding CH_4_ production in freshwater. Phosphonate compounds were once thought to be resistant to breakdown and their formation remains poorly understood (17, 65). Under P limiting conditions, competition for these and other P compounds among multiple groups of organisms will be intense, and potentially variable in space and time. Along with variations in CH_4_ oxidation relative to production rates, this can drive variations in CH_4_ production such as those evident in our data.

As a newly-identified CH_4_ source that occurs via an unknown pathway (23), we used a combination of approaches to identify whether photosynthetic CH_4_ production occurs in freshwater, and by what means. Although independent data types (CH_4_ concentrations, treatment effects, isotopic data, ‘omic data) may be interpreted in multiple ways, the combined data are consistent with photosynthetic production in the UC4 experiment (Figs. 3 and 4). For example, the high light and BES treatments boosted CH_4_ concentrations (Fig. S2), but this CH_4_ is unlikely to have been the result of methanogenesis or MPn breakdown given low and decreasing *mcrA* and *phnJ* expression (Fig. 4). Instead, COR and DPOR transcripts were present and responsive to treatments as shown by metagenomic and metatranscriptomic data. Our data are further indicative of an isotopically light CH_4_ source (Fig. 3), which is inconsistent with production from MPn (44, 45). Extending from the UC4 experiment, photosynthetic CH_4_ production may take place in at least five other experiments where BES and high light also significantly increased CH_4_ (L2, L3, LG4, UC3, UC4; Fig. S2). Dark treatments further suggest photosynthetic production could occur in several additional experiments alongside increased CH_4_ oxidation (Fig. S2). Finally, the ubiquity of COR and DPOR genes and transcripts indicates that the potential for photosynthetic CH_4_ production is widespread (Fig. 5), yet the apparent variability across experiments is consistent with known variability in photosynthesis by *Limnohabitans* (51, 66).

These patterns were most clearly evident in experimental treatments, highlighting the efficacy of including different treatments to examine paradoxical CH_4_ production. We leveraged data from these treatments to identify potential CH_4_ production mechanisms—providing an experimental approach, isotopic signature data, and two potential gene targets to examine in other aquatic ecosystems. We found that this mechanism is not limited to the Cyanobacteria, as the majority of DPOR and COR transcripts and genes were derived from the betaproteobacteria. This has several important implications. First, *Limnohabitans* and *Polynucleobacter* are among the most abundant and ubiquitous bacteria found in freshwater ecosystems (31). We recovered MAGs from both of these groups, and all contained DPOR and COR genes, as well as genes for phosphonate metabolism. *Limnohabitans* also expressed *phnJ* genes, indicating that they can play a dual role in paradoxical CH_4_ production. These findings demonstrate that two mechanisms for paradoxical CH_4_ production co-occur in multiple lineages of abundant and widespread bacteria in freshwater. Second, our data are indicative of an additional paradoxical CH_4_ production mechanism: CH_4_ production by aerobic anoxygenic photosynthetic (AAnP) bacteria. AAnP bacteria are also prevalent in the ocean, where they constitute up to 10% of bacterial communities and play an important role in ocean carbon cycling (66). Given their significance, the potential for CH_4_ production by marine AAnP deserves examination. More broadly, COR and DPOR are nitrogenase-like enzymes that have a central role in photosynthesis. Multiple other nitrogenase-like enzymes produce CH_4_ (62, 64), and our results implicate COR and DPOR. However, this mechanism is variable across our experiments. This raises the possibility that the potential for photosynthetic CH_4_ production is widespread, but not always active. Understanding the factors that regulate photosynthetic CH_4_ production (such as light, compounds like BES, and other factors) is therefore important. Our combined dataset consequently demonstrates that paradoxical CH_4_ production is complex, with several contributing processes—some present within the same organisms—each affected by a range of environmental variables (e.g., P availability, light), and all producing CH_4_ to differing degrees. Understanding this complexity is essential, as multiple paradoxical CH_4_ production mechanisms are widely-distributed in aquatic ecosystems, where they may represent important sources of CH_4_ to the atmosphere.

## Materials and Methods

### Field site, sample collection, and experimental set up

Water samples were collected in 2016-2018 in five high-elevation lakes in Yosemite National Park, and used in incubation experiments. Samples were collected in Lukens (L), Lower Cathedral (LC), Upper Cathedral (UC), Lower Gaylor (LG), and Upper Gaylor (UG) Lakes. Experiments are denoted by lake abbreviation and sequential numbering (Table S5). Water samples were collected in the littoral and limnetic zones of the lakes at 0.1 m depth with acid-washed cubitainers, and then kept on ice or refrigeration to maintain in-lake temperature until laboratory incubations were established within 24 hours of sample collection. Temperature and dissolved oxygen were measured at the time of sample collection using a ProODO YSI probe (YSI Inc., Yellow Springs, OH, USA).

Water collected in the lakes was transferred to 170 or 300 ml Wheaton bottles, capped and crimped, and a known volume of air was introduced to generate a headspace for sampling. Initial CH_4_ samples were collected, and bottles were incubated in water baths at constant temperature. All incubations were run in triplicate and set to the temperature at which water samples were collected in the field. The different treatments used in the experiments were:

- Control: unamended lake water following a natural day-night set up (water bath lid was opened at 7:00 hours and closed at 18:00 hours).
- Dark: unamended lake water; bottles were kept in the dark during the whole incubation time.
- BES: lake water was amended with 2-bromoethanesulphonate (BES) to a final concentration of 5×10^-4^ M. This concentration has been established to inhibit methanogens (25) and has been used in methane paradox experimentation before (8). This treatment followed the same natural daylight set up as the control.
- High light: unamended lake water was subjected to 500 mmol/m^2^ inside a growth chamber and followed the same natural daylight set up as the control (light would turn on at 7:00 hours and off at 18:00 hours).
- Sterile treatments: 0.22 μm-filtered lake water did not show significant CH_4_ production or oxidation over time, indicating observed CH_4_ production and consumption is likely biotic.

For all incubation types, gas samples were taken from the headspace every 6-24 hours for up to 96 hours with a syringe and immediately transferred to exetainers for later analyses in a gas chromatograph. Not all treatments were tested in each incubation experiment. Temperature and oxygen concentrations were monitored at each sampling point. Optical sensor spots (Fibox, Loligo Systems, Viborg, Denmark) were used to measure oxygen concentrations during incubations (detection limit of 100 nM) and make sure that the water did not go anoxic at any time. Temperature was measured with a Fibox temperature sensor and kept constant during the incubation time. Water samples were filtered at the beginning and end of the incubation for DNA and RNA sampling.

### Methane measurements

Methane concentrations were measured via headspace equilibration and gas chromatography. Headspace gas samples from incubations were collected with a gas-tight syringe into 12-mL Labco Exetainer vials (Labco Ltd., Lampeter, Ceredigion, UK) after incubations bottles were shaken for 2 minutes to reach equilibration. Samples were subsequently analyzed using a Shimadzu GC-2014 gas chromatograph with flame ionization detection (FID) for CH_4_ (67). Headspace CH_4_ concentration measurements were then used to calculate CH_4_ concentration in lake water based on Henry’s law of equilibrium (68).

### DNA and RNA Extraction

Water samples were filtered through 0.22 μm (Millipore, Darmstadt, Germany) then DNA filter samples were preserved in Sucrose-Tris-EDTA (STE) buffer in pre-prepped Lysis Matrix E tubes and frozen at −80°C until extraction. RNA samples were preserved in RNA*later*^®^ (Ambion™, AM7021) in pre-prepped Lysing Matrix E tubes (MP Bio, Eschwege, Germany), and frozen at −80°C until extraction. DNA was extracted using the Qiagen DNeasy Blood & Tissue Kit with a modified protocol from Beman et al. (69). Briefly, samples were lysed with 100μL 10% sodium dodecyl sulfate (SDS), DNA gets separated from proteins and cellular debris using proteinase K (20mg mL^-1^; Qiagen, Inc., Valencia, CA, USA), precipitated with ethanol and cleaned up. After extraction samples were preserved at −80°C until further analyses. RNA was extracted using a *mir*Vana miRNA Isolation Kit (Ambion™, AM1560) with a modified protocol from Huber and Fortunato (70). Briefly, samples are lysed with the kit’s lysing matrix, then subjected to an organic extraction with phenol chloroform followed by a wash to obtain RNA. Immediately after RNA extraction we used the SuperScript III First-Strand Synthesis System for RT-PCR (Life Technologies Corporation, Carlsbad, CA, USA) to synthesize first-strand cDNA and samples were preserved at −80°C until further analyses. DNA, RNA and cDNA purity was measured using a Biospectrometer (Eppendorf AG, Hamburg, Germany) and the concentrations were quantified using PicoGreen Quant-iT dsDNA quantitation assay (ThermoFisher Scientific, USA) for DNA samples and the MaestroNano Pro (Maestrogen Inc., Taiwan) for RNA and cDNA samples.

### 16S sequencing

DNA and RNA extracted from filtered water samples were diluted to a common concentration (1ng/ul) and sent for 16S rRNA amplicon sequencing on an Illumina MiSeq (Illumina, San Diego, CA, USA) according to Earth Microbiome protocols. We used the universal primers 515F-Y (5’-GTGYCAGCMGCCGCGGTAA) and 926R (5’-CCGYCAATTYMTTTRAGTTT). DNA samples were sequenced at the Joint Genome Institute (Berkeley, CA, USA) and RNA samples at the Argonne National Laboratory (Lemont, IL, USA).

ASVs were generated from 16S rDNA and rRNA sequence data using the Divisive Amplicon Denoising Algorithm (DADA2; 71) as implemented in QIIME 2 (72), and then used for subsequent analyses. After import and demultiplexing, read quality was visualized using the ‘qiime tools view’ command. Reads were then processed using the ‘qiime dada2 denoise-paired’ command, with 13 bp trimmed from both the forward and reverse reads, truncation of reverse to 169 bp (due to the well-known decline in sequence quality observed for MiSeq reverse reads), and training of the denoising algorithm on 1 million reads. Classification of ASVs was conducted in mothur (73) using the SILVA (version 128) database.

### Metatranscriptomics and metagenomics

Metatranscriptomes and metagenomes were generated from later experiments in order to examine potential production mechanisms and coupled methane oxidation. Following extraction, DNA and RNA samples were sent for metagenome/metatranscriptome sequencing in the Vincent J. Coates Genome Sequencing Laboratory (GSL) at the University of California, Berkeley (https://genomics.qb3.berkeley.edu/), which is supported by NIH S10 OD018174 Instrumentation Grant. For each DNA sample, 250 ng of genomic DNA was sheared and libraries were prepared using the KAPA HyperPrep Kit (Kapa Biosystems, Wilmington, MA, USA). For each RNA sample, ~800 ng of total RNA was depleted of rRNA using the Ribo-Zero rRNA Removal Kit (Illumina, Inc., San Diego, CA, USA), sheared, and libraries were prepared using the KAPA RNA HyperPrep Kit (Kapa Biosystems, Wilmington, MA, USA). 12 samples were pooled into a single lane and sequenced via 150-cycle paired-end sequencing on the Illumina HiSeq 4000 platform (Illumina, Inc., San Diego, CA, USA).

Data were demultiplexed by the GSL and reads were filtered and trimmed using BBDuk (https://jgi.doe.gov/data-and-tools/bbtools/bb-tools-user-guide/bbduk-guide/) with the following parameters: maq=8, maxns=1, minlen=40,minlenfraction=0.6, k=23, hdist=1, trimq=12, qtrim=rl. Forward and reverse reads were then merged using PANDASeq (https://github.com/neufeld/pandaseq; 74) with default parameters. Merged reads were subsequently annotated in DIAMOND (http://diamondsearch.org/; 75) using the NCBI NR database (accessed February 11^th^, 2020) with the following search criteria: maximum number of target sequences = 1, bit-score > 40. In order to quantify functional gene abundances, we filtered the DIAMOND annotation data to find the number of functional genes of interest present in each metagenome or metatrascriptome, discarding those with similarity below 60%. We calculated percent abundance based on the total number of reads and the number of targeted genes in each metagenome or metatranscriptome. Metagenomic contigs were assembled via megahit v 1.1.3 (76) using an initial k-mer size of 23, and were annotated as described above but with the ‘long reads’ option in DIAMOND.

For additional analyses, UC4 metatranscriptomes were uploaded to the Joint Genome Institute (JGI; https://img.jgi.doe.gov/) Integrated Microbial Genomes & Microbiomes platform for additional analyses. We used the function comparisons tool to identify functions that were significantly different in the treatments vs. the control, and the function category tool identify differences in KEGG pathway categories among the treatments. Moreover, we used the phylogenetic distribution tool to quantify the abundance of different phylogenetic groups of potential importance in the methane paradox. Metatranscriptome reads were also mapped to MAGs using bowtie2 v 2.3.4.3 (77) and visualized using Anvi’o (78). Coverage files were generated in samtools v 1.11 (http://www.htslib.org/) for statistical comparisons. MAGs were assembled as described by Nayfach et al. (53).

### Stable isotopic measurements

Headspace of incubation bottles was transferred to evacuated exetainers and sent to the UC Davis Stable Isotope Facility (Davis, CA) for analysis. Measurements of stable isotope ratios of carbon (δ^13^C) in CH_4_ were conducted using a ThermoScientific Precon concentration unit interfaced to a ThermoScientific Delta V Plus isotope ratio mass spectrometer (ThermoScientific, Bremen, Germany). In brief, gas samples are passed through a H_2_O/CO_2_ scrubber and a cold trap, and CH_4_ is then separated from other gases and oxidized to CO_2_. Pure CO_2_ reference gas is used to calculate provisional δ values, and final δ values are calculated after correcting for changes in linearity and instrumental drift, and expressed relative to the Vienna PeeDee Belemnite (V-PDB) standard.

### Statistical analyses

Spearman’s correlation test was used to assess the relationship between time (h) and CH_4_ concentration in all incubations and across all treatments. A priori significance level was defined as α<0.05. A significantly positive relationship between time and CH_4_ was considered as net CH_4_ production in the incubation, whereas a significantly negative relationship was considered as net CH_4_ consumption; otherwise we defined the incubation as not having significant net production or consumption over time. We subsequently used ANOVA and Tukey honestly significant difference (HSD) post-hoc tests to test for cases where CH_4_ production or oxidation varied significantly between sampling time points, resulting in nonlinear patterns in CH_4_ concentrations over the course of the experiment. For example, initial CH_4_ production could be followed by subsequent oxidation, resulting in significant CH_4_ increases followed by significant decreases.

To express the different responses in terms of CH_4_ concentration over time among the different experiments treatments we first calculated the response ratios (lnRR) by the following equation:

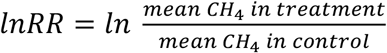

Additionally, we calculated which treatments were significantly different from each other and the control using ANOVA and Tukey honestly significant difference (HSD) post-hoc tests. All statistical analyses and figures were done in the R statistical environment (RStudioVersion 1.2.5001).

## Acknowledgments and funding sources

This work was supported by the University of California through the Valentine Eastern Sierra Reserve graduate student research grant, the Institute for the Study of Ecological and Evolutionary Climate Impacts fellowship, the University of California Merced Environmental Systems summer grants, and the UC MEXUS-CONACYT doctoral fellowship. We thank the United States National Park Service for facilitating sample collection in Yosemite National Park under permits YOSE-2016-SCI-0118, YOSE-2017-SCI-0104 and YOSE-2018-SCI-0091. We thank Steve Hart for his help processing methane samples, Jay Sexton for his help with incubation chambers, and Angela Yu, Joaquin Fraga, Sonia Vargas, Ariadna Cairo, Samantha Vazquez, Jorge Montiel and Daniela Alonso for their help with field and laboratory work.

## Supplementary Materials for

### Supplementary Information Text

#### Methane production and consumption over time

All experiments were conducted for at least 24 hours to permit direct comparisons across experiments. However, we included multiple sampling time points before 24 hours in some experiments, while others were conducted for up to 96 hours (Fig. S1). Although longer incubations may introduce bottle effects, this has been used in earlier work (1, 2), and provides multiple comparisons through time, between treatments, and across experiments.

In experiments with multiple sampling time points before 24 hours, we occasionally observed initial decreases that were followed by increases by 24 hours (Fig. S1). This could be explained by the development of P limitation that triggers methane (CH_4_) production; for example, Karl et al. (3) observed CH_4_ production by MPn breakdown only under P-stressed conditions. Other incubations showed the opposite pattern: an initial increase followed by a decrease in concentration (L2, UC2, LG1). Longer experiments likewise showed eventual decreases in CH_4_ concentrations after 24 hours (e.g., L4, L7, LC1, UC4). This behavior has been observed in earlier studies (4) and was the intention of these experiments; specifically, longer incubations may capture the time lag between initial production followed by eventual consumption once CH_4_ concentrations exceed a required threshold for methanotrophy. Put another way, both CH_4_ production and consumption are taking place, but production rates exceed oxidation rates until oxidation ‘catches up’ and eventually exceeds production. In general,*pmoA* transcripts were more abundant when CH_4_ concentrations in the incubations were higher (L7, LG5 and LG6; CH_4_ >100 nM)—consistent with threshold-dependent CH_4_ oxidation (4). A similar pattern was evident in metagenomes, as *pmoA* genes were absent in the LG2 incubations where the CH_4_ concentrations were <50 nM, and in low abundance in the L6 incubations (CH_4_ = ~100 nM).

**Fig. S1.**
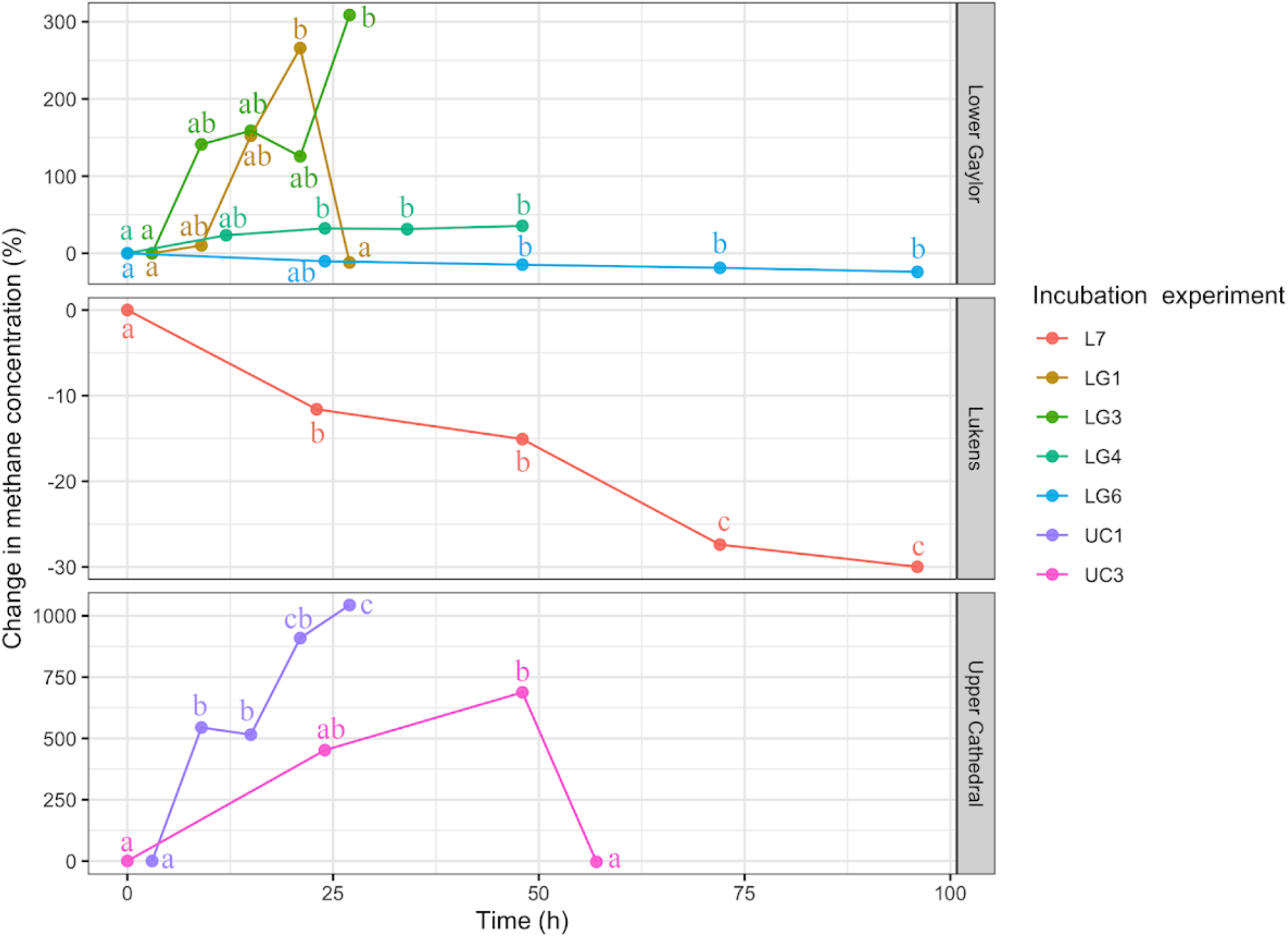
Percent change in methane (CH_4_) concentrations over time. Different colors represent each incubation experiment with statistically significant differences in CH_4_ concentrations among different time points. Letters represent similarities or differences between time points according to Tukey HSD test.

**Fig. S2.**
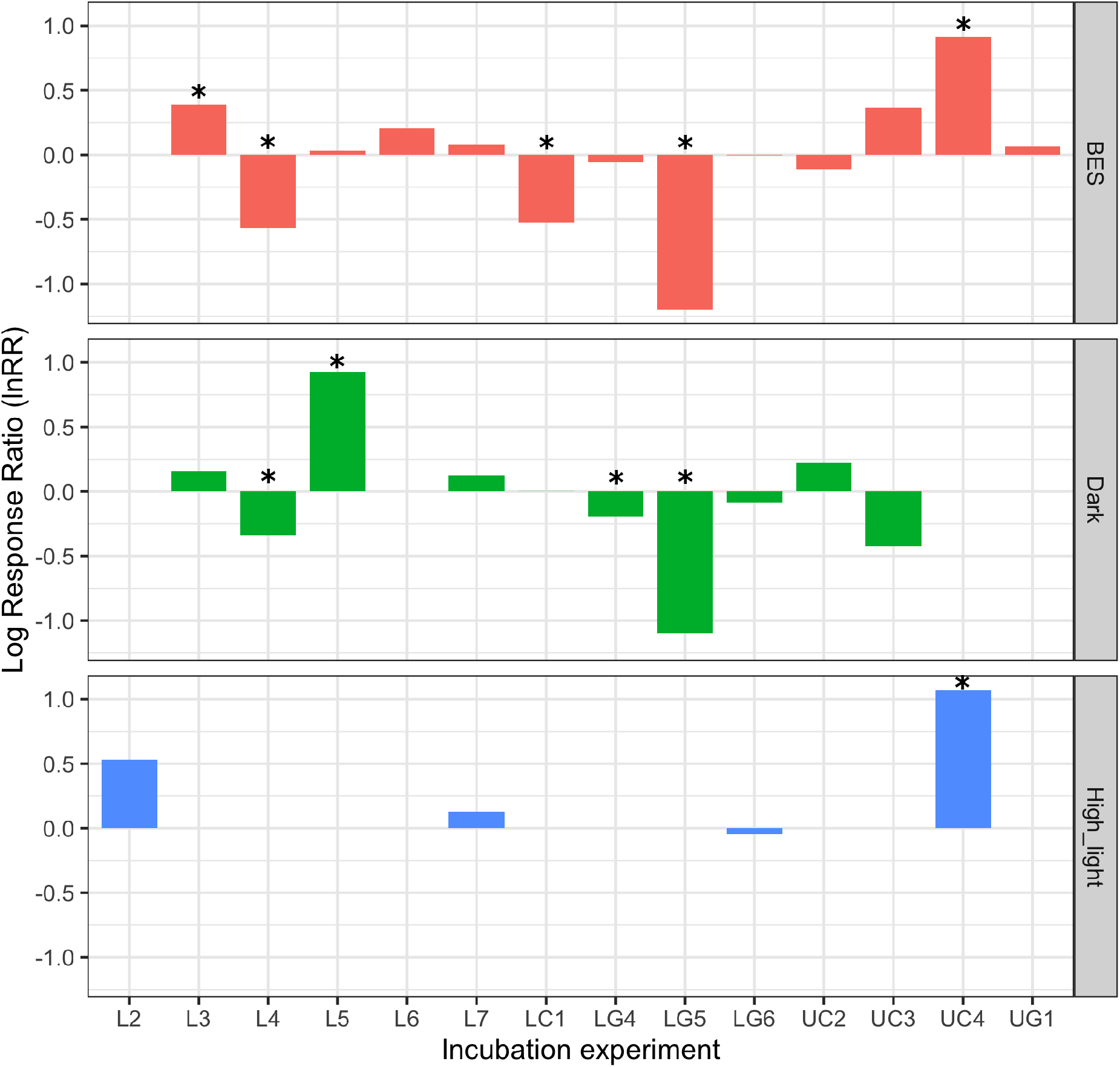
Log response ratios for methane production rates in experimental treatments. Asterisks denote experimental treatments that were significantly different from the control. Experiments L3, L5, LG5 and UC3 were longer experiments and treatment effects are shown for 36, 74, 86 and 57 hours respectively (all others are shown at 24 hours).

**Table S1.** Percentage of *phnJ*, DPOR, and COR transcripts and genes, as well as the mean and range of percent identity (ID%), for significant microbial taxa in metatranscriptomes and metagenomes. Blank cells indicate that a particular group is irrelevant for that particular gene.

|  | *Comamonadaceae* | Cyanobacteria | *Polynucleobacter* | Other<br>*Burkholderiales* |
| --- | --- | --- | --- | --- |
| % of *phnJ* transcripts | 20.9 |  |  |  |
| *phnJ* transcripts mean (and range) ID% | 92.2 (69.6-100) |  |  |  |
| % of *phnJ* genes | 27.4 |  |  |  |
| *phnJ* genes mean (and range) ID% | 84.8 (60.4-100) |  |  |  |
| % of DPOR transcripts | 30.6 | 6.3 | 5.7 | 2.8 |
| DPOR transcripts mean (and range) ID% | 88.8 (60.4-100) | 91.5 (63.5-100) | 94.5 (69.7-100) | 92.1 (60.2-100) |
| % of DPOR genes | 25.7 | 0.3 | 35.3 | 1.4 |
| DPOR genes mean (and range) ID% | 86.2 (60-100) | 86.9 (64.4-100) | 89.8 (60-100) | 88.2 (60-100) |
| % of COR transcripts | 48.7 |  | 5.8 | 6.8 |
| COR transcripts mean (and range) ID% | 91.5 (60-100) |  | 94.3 (61.4-100) | 93.6 (61.7-100) |
| % of COR genes | 29.8 |  | 34.1 | 2.4 |
| COR genes mean (and range) ID% | 88.5 (60-100) |  | 90.7 (60-100) | 87.3 (60-100) |

**Table S2.** Percent identify of 16S rRNA sequences from abundant *Comamonadaceae* ASVs to those in sequenced genomes in the RefSeq Genome Database.

| ASV | %ID | Genome |
| --- | --- | --- |
| b66e24bf473f06a1fc27fa300549636c | 99.7 | Limnohabitans Rim47 and others, Acidovorax radialis and others |
| 14f5d35b60a4aa29fb4630dc0531b12e | 99.1 | Acidovorax temperans, delafeldii and other strains |
| f361f2ed2d8297ddacd4e36526eb8265 | 100 | Limnohabitans Rim11 |
| f2fce6a8c3a54fda2ae66977fd9b45af | 99 | Limnohabitans Rim11 and MMS10 |
| 222b12c7663803b96339c5af33e2b3ae | 100 | Limnohabitans 103DPR2 |
| 9b6d5192b8dc30e78ce626298c8a0a26 | 99.4 | Limnohabitans Rim47 and others, Acidovorax radialis and others |
| 95dd5f1247bb97a138d1fa91e409f85 | 99.3 | Limnohabitans 103DPR2 and JirII |
| 62767018de03c34a4305384df06ce393 | 98.8 | Acidovorax delafeldii, others |
| 86832a3826a81807da70809f71eed601 | 99.4 | Limnohabitans Rim47 and others, Acidovorax radialis and others |
| 5efb46c72256df114e8f2b354fb44192 | 99.7 | Limnohabitans Rim11 and MMS10 |
| d81e17e704250a1f80e1005be449b274 | 99.4 | Limnohabitans 103DPR2 and JirII |
| 12e95e1ebf7046141228b71394bd745b | 99.7 | Limnohabitans planktonicus, Rim28, and others |
| 02523daf5356885c7d28e05d5910acb3 | 100 | Rhodoferrax sediminis, ferrireducens, saidenbachensis, others |
| 8f8863b1fba0419252ce1756ec2f9b46 | 100 | Rhodoferrax bucti strain GSA243-2 |
| 1b24e60fca2c1ce5d0e34cab586c4537 | 100 | Polaromonas sp. YR568 |
| d20e921d6be99c2d20fe3d688f878774 | 99.7 | Rhodoferrax bucti strain GSA243-2 |
| 74dace2780cac91c5a0d5598db981acc | 100 | Limnohabitans Rim47 and others, Acidovorax radialis and others |
| 6f6aa478c9f6ad0d6f8b0a415706fa81 | 100 | Limnohabitans curvus, others |
| 317124a0271aaea477061ada50e0cfb | 99.4 | Limnohabitans Rim8, others |
| c4143240980a12d00bf11daf4fefbd10 | 99.7 | Limnohabitans Rim8, others |
| 12fd09c14da54c5c7c412f7e493c4e16 | 100 | Limnohabitans planktonicus, others |
| c4d004171ee776f1e529ea5dcb010508 | 100 | Rhodoferrax koreense strain DCY-110, Curvibacter sp. AEP1-3 |
| 405bafec3faf0d7ba859b721abc30c01 | 98.8 | Comamonas terrigena (multiple strains) |
| 1f796798ba249fd048e233237b86bf05 | 98.8 | Pelomonas (multiple) |

**Table S3.** Examples of multiple*phn* genes co-located on assembled contigs (>10,000 base pairs in length) within metagenomes from the LG2 and L6 experiments.

| Exp. | Contig | bp | *phnV* | *phnO* | *phnN* | *phnM* | *phnL* | *phnK* | *phnJ* | *phnI* | *phnH* | *phnG* | *phnF* | *phnE* | *phnD* | *phnC* |
| --- | --- | --- | --- | --- | --- | --- | --- | --- | --- | --- | --- | --- | --- | --- | --- | --- |
| LG2 | k141_33043 | 23126 |  | 1 | 1 | 1 | 1 | 1 | 1 | 1 |  |  |  | 1 | 1 | 1 |
| LG2 | k141_67057 | 20078 |  |  | 1 | 1 | 1 | 1 | 1 | 1 | 1 | 1 |  |  |  | 1 |
| LG2 | k141_174869 | 50613 |  |  | 1 | 1 | 1 | 1 | 1 | 1 | 1 | 1 | 1 | 1 |  |  |
| LG2 | k141_61413 | 94539 |  |  | 1 |  | 1 | 1 | 1 | 1 | 1 | 1 |  | 1 |  | 1 |
| LG2 | k141_68876 | 34960 |  |  | 1 | 1 | 1 | 1 | 1 | 1 | 1 | 1 | 1 | 1 |  | 1 |
| L6 | k141_232042 | 14329 | 1 |  |  |  |  |  | 1 | 1 | 1 | 1 |  |  |  |  |
| L6 | k141_58845 | 164079 | 1 |  |  |  | 1 | 1 | 1 | 1 | 1 | 1 |  | 1 |  |  |
| L6 | k141_137475 | 51136 |  |  | 1 |  | 1 | 1 | 1 | 1 | 1 | 1 |  |  | 1 | 1 |

**Table S4.** List of betaproteobacterial metagenome-assembled genomes (MAGs) and links to scaffolds containing COR, DPOR, and phosphonate metabolism genes on the Integrated Microbial Genomes Microbiomes system.

| MAG | Affiliation | COR gene scaffold(s) | DPOR gene scaffold(s) | Phosphonate metabolism genes scaffold(s) |
| --- | --- | --- | --- | --- |
| 1 | *Rubrivivax* | Ga0136641_1000428 | Ga0136641_1000214 | Ga0136641_1000005<br>Ga0136641_1000835 |
| 5 | *Limnohabitans* | Ga0136641_1000705 | Ga0136641_1000769<br>Ga0136641_1001408 | Ga0136641_1001116<br>Ga0136641_1001732 |
| 6 | *Limnohabitans* | Ga0136641_1000147 | Ga0136641_1000390<br>Ga0136641_1003698 | Ga0136641_1000400<br>Ga0136641_1004149 |
| 18 | *Limnohabitans* | Ga0136642_1003267 |  | Ga0136642_1004820 |
| 20 | *Polynucleobacter* | Ga0136642_1002146 | Ga0136642_1003437 |  |

**Table S5.** Details of experimental incubations, including: experiment number; year, month, and days of experiment; whether methane production was significant and P-value; generation of 16S rDNA, 16S rRNA, metatranscriptomes, and metagenomes; length of experiment; and treatments. BES corresponds to the addition of 2-bromoethanesulphonate as a methanogenesis inhibitor, Dark represents experiments incubated in total darkness and HL (high-light) are experiments subjected to high-light intensity during the incubation.

| Exp. | Y | M | D | CH<sub>4</sub> | P | 16S rDNA | 16S rRNA | Metat | Metag | Length (hr) | Treatments |
| --- | --- | --- | --- | --- | --- | --- | --- | --- | --- | --- | --- |
| L1 | 2016 | 8 | 12-13 | + | 0.02 | X |  |  |  | 24 |  |
| L2 | 2016 | 9 | 11-12 | + | 0.02 | X |  |  |  | 24 | HL |
| L3 | 2017 | 7 | 28-30 |  |  | X |  |  |  | 48 | BES, Dark |
| L4 | 2017 | 8 | 8-10 |  |  | X |  |  |  | 48 | BES, Dark |
| L5 | 2017 | 9 | 26-29 | - | 0.02 |  | X |  |  | 74 | BES, Dark |
| L6 | 2017 | 11 | 13-15 |  |  | X | X | X | X | 55 | BES |
| L7 | 2018 | 9 | 23-28 | - | 0.02 |  | X | X |  | 96 | BES, Dark, HL |
| LG1 | 2016 | 8 | 8-9 | +/- | 0.02 | X |  |  |  | 24 |  |
| LG2 | 2016 | 9 | 9-10 |  |  | X |  |  | X | 24 |  |
| LG3 | 2016 | 10 | 7-8 | + | 0.02 | X |  |  |  | 24 |  |
| LG4 | 2017 | 8 | 24-26 | + | 0.02 | X |  |  |  | 48 | BES, Dark |
| LG5 | 2017 | 10 | 1-4 |  |  |  | X | X |  | 86 | BES, Dark |
| LG6 | 2018 | 9 | 10-14 |  |  |  | X | X |  | 96 | BES, Dark, HL |
| UC1 | 2016 | 8 | 10-11 | + | 0.02 | X |  |  |  | 24 |  |
| UC2 | 2017 | 8 | 10-12 |  |  |  |  |  |  | 48 | BES, Dark |
| UC3 | 2017 | 10 | 8-11 | +/- | 0.006 |  | X |  |  | 57 | BES, Dark |
| UC4 | 2018 | 10 | 8-12 |  |  |  | X | X |  | 96 | BES, Dark, HL |
| LC1 | 2017 | 08 | 26-28 | - | 0.02 |  |  |  |  | 48 | BES, Dark |
| UG1 | 2017 | 08 | 12-14 |  |  |  |  |  |  | 48 | BES |

